# Divergent Neurobehavioral Effects of CFTR Modulators Elexacaftor and Ivacaftor in Mice

**DOI:** 10.1101/2025.07.02.662194

**Authors:** Qian Ge, Amy Fagan, Hui Lu, Jianyang Du, Leah R. Reznikov

## Abstract

Recent advances in cystic fibrosis transmembrane conductance regulator (CFTR) modulator combination therapies have markedly improved survival and quality of life for people with cystic fibrosis (CF). Among these, the elexacaftor-tezacaftor-ivacaftor combination (Trikafta) and its updated formulation, vanzacaftor-tezacaftor-deutivacaftor (Alyftrek), represent major therapeutic milestones. However, emerging reports suggest that Trikafta may contribute to anxiety and depression in some people with CF. In this study, we investigated the neurobehavioral effects of elexacaftor and ivacaftor in mice. Acute administration of elexacaftor elicited anxiety-like behavior, while ivacaftor induced depressive-like behavior. Additionally, we confirmed the presence of *Cftr* mRNA in the amygdala and hippocampus, brain regions implicated in anxiety and depression. These findings provide preclinical support for the reported mental health side effects experienced by some people with CF and offer new insights into CFTR function within the central nervous system, potentially guiding the development of future CF therapies with improved neuropsychiatric profiles.

## Introduction

Cystic fibrosis (CF) is a life-shortening, autosomal recessive disease caused by mutations in the *CFTR* gene, which encodes the cystic fibrosis transmembrane conductance regulator (CFTR) protein. CFTR is an anion channel responsible for transporting chloride and bicarbonate across epithelial cell membranes. Approximately 1,000 new cases of CF are diagnosed each year. Over the past decade, a new class of therapies known as CFTR modulators has dramatically improved outcomes for people with CF by targeting the underlying protein defect (1). In 2012, ivacaftor became the first CFTR potentiator approved for clinical use, enhancing the function of CFTR channels at the cell surface (2). This was followed in 2015 by the combination therapy lumacaftor/ivacaftor, which added a CFTR corrector to improve protein folding and trafficking (3). In 2019, the introduction of Trikafta (elexacaftor/tezacaftor/ivacaftor) marked a major advancement, offering the first triple combination therapy that significantly improved lung function and quality of life for a broader population of people with CF (2). These modulators work by either potentiating CFTR channel gating (ivacaftor) or correcting CFTR folding and trafficking (lumacaftor, tezacaftor, elexacaftor).

Some people with CF experience depression and anxiety (4). For instance, one report suggested that more than 35% of people with CF in the United States have depressive symptoms (5). Depression in people with CF has been associated with poorer lung function (6), as well as increased CF-related healthcare costs (7). Some reports also suggest that people with CF have roughly double the rates of anxiety compared to the general population (8, 9), leading to poorer medical outcomes (7-9). In studies involving Trikafta, worsening or onset of depression or anxiety has been reported as an adverse side effect (10). Despite these observations, potential mechanisms linking Trikafta to worsening mental health in some individuals are poorly understood.

While lack of anion transport, notably chloride and bicarbonate, across epithelia is accepted as the basic defect in CF, it is well established that CFTR is expressed in the nervous system (11). The expression of CFTR in the nervous system is intriguing, given the central role of chloride in regulating neuronal excitability. Chloride homeostasis is a fundamental mechanism underlying inhibitory neurotransmission, and many pharmacological agents exert their effects by modulating this balance. Among the most well-known are the benzodiazepines. Benzodiazepines serve as clinical anxiolytics that potentiate the activity of GABA A receptors (12). Activation of GABA A receptors typically results in chloride influx into a neuron (13), leading to neuronal inhibition. This raises an intriguing possibility that modulation of CFTR in the nervous system may have neurobehavioral effects that are not yet fully understood. If true, this mechanism may offer valuable insight into the reported adverse mental health effects in some people with CF prescribed Trikafta.

In the current study, we investigated whether an acute intraperitoneal injection of elexacaftor or ivacaftor modulated neurobehavioral outcomes in mice, with an emphasis on anxiety-like and depressive-like behaviors. Additionally, we confirmed the presence of *Cftr* mRNA in mouse brain regions involved in regulating emotional behaviors, suggesting a potential physiological mechanism linking elexacaftor and ivacaftor to neurobehavioral outcomes.

## Materials and methods

### Mice

Specific pathogen-free 6-9 weeks male and female C57BL/6 mice were purchased from the Jackson Laboratory. To ensure scientific rigor, experimental factors such as sex, age, and weight were closely matched throughout the study. Experimental mice were maintained on a standard 12-hour light-dark cycle and received standard chow and water *ad libitum*. Animal care and procedures complied with National Institutes of Health standards. All experimental protocols were approved by the Institutional Animal Care and Use Committees (IACUC) of the University of Tennessee Health Science Center (UTHSC), protocol number 24-2696. The number of animals used in each experiment was determined based on power calculations to ensure statistical significance while adhering to the principles of reduction, refinement, and replacement. These calculations considered expected effect size, variability, and desired statistical power.

### Behavioral experiments

We conducted well-established behavioral tests in mice as indicated below. For all behavioral tests, mice were acclimated to the testing room for ≥ 30 minutes to minimize stress from novel environments. Mice were gently handled for 3-5 minutes daily for 3 days to reduce handling-related stress. More details about specific tests are shown below.

### Anxiety-like behaviors

#### Open field test (OFT)

We assessed exploratory and anxiety-like behaviors in an open field box for 6 minutes. The OFT was used to determine the general activity levels, gross locomotor activity, exploration habits of mice, and anxiety-like behaviors. The evaluation occurred within a cubic, white plastic enclosure measuring 30 cm × 30 cm × 30 cm. Mice were introduced to a corner of the arena, given 5 minutes to acclimate to the unfamiliar surroundings, and recorded via video. Subsequently, the recorded footage, encompassing metrics such as distance traversed, velocity, resting intervals, movement duration, and time spent in both the center and corners of the arena, was analyzed using Anymaze software. This examination serves as an initial assessment of locomotor activity and anxiety-related behaviors, particularly focusing on the duration spent in the central area.

#### Elevated plus maze test (EPM)

The EPM is a gold-standard behavioral assay to evaluate anxiety-like behavior in rodents. It exploits the innate conflict between a mouse’s curiosity to explore novel environments and its aversion to open, elevated spaces. Reduced time spent in open arms and fewer open-arm entries are interpreted as heightened anxiety. We evaluated anxiety-like behavior by recording entries into and time spent in the open arms of a maze for 6 minutes. It was conducted as previously described (14). The EPM apparatus, 40 cm high from the floor, consists of two open arms (35 × 10 cm) and two closed arms (35 × 10 cm), those two parts stretch perpendicular to each other and connect to a central platform (5 cm). Mice were placed in the center zone facing an open arm and allowed to explore the maze freely for 5 minutes. The time spent in the open arms was measured, and the percentage of open-arm time was calculated by dividing the time spent in the open arms by the total time spent in both the open and closed arms (5 minutes). Entries into the open arms were analyzed by calculating the percentage of open-arm entries relative to the total number of entries into both the open and closed arms.

#### Marble burying test (MBT)

MBT is a mouse model of anxiety-like behavior in mice. It is based on the observation that mice bury either harmful or harmless objects in their bedding when feeling anxious. In this paradigm, mice are placed in a transparent cage containing a 5-cm layer of bedding material, with 16 marbles (1.5 cm diameter) arranged in a 4×4 grid. Mice are acclimated to the testing room for ≥ 1 hour before the experiment. Individual mice are placed gently into the center of the cage and are allowed to explore the cage for 10 min under standardized conditions, including dim lighting (∼50 lux) and background white noise (55-65 dB). The apparatus is cleaned with 70% ethanol between trials to eliminate odor cues. We record the number of marbles buried (defined as ≥ 2/3 covered by bedding), with higher marble burial typically interpreted as increased anxiety.

### Depression-like behaviors

#### Novelty suppressed feeding test (NSFT)

The use of the NSFT, in which exposure to a novel environment suppresses feeding behavior, has been used to assess chronic depression-related behavior in animals. Mice are food-deprived for 24 hours (water *ad libitum*). During the test phase, a single food pellet (highly palatable food, that was pre-exposed to the mice before food deprivation) was placed centrally on a small dish (5 cm diameter) in the center of a novel open-field arena (50 cm × 50 cm × 40 cm). Mice were placed in a corner of the novel arena. We recorded behavior for 10 minutes using a camera placed over the open field and analyzed the video with Anymaze software. The latency to initiate feeding (time until the mouse takes the first bite) was quantified in a 10-minute session. We cleaned the arena with 70% ethanol between trials to eliminate food odor cues. Prolonged latency was regarded as correlating with heightened depression-like states.

#### Tail suspension test (TST)

The TST is a widely used behavioral assay to evaluate depression-like states in rodents, particularly “behavioral despair” or learned helplessness. Mice suspended by their tails initially struggle to escape but eventually adopt a posture of immobility, which is interpreted as a resignation to stress. A sound-attenuated testing chamber (30 × 30 × 40 cm) was constructed from an opaque acrylic to prevent visual distractions. A horizontal bar was positioned 30-50 cm above the chamber floor, allowing mice to hang freely without touching surfaces. Adhesive tape is used to suspend mice by the tail, ensuring secure attachment while minimizing tissue damage. Mice are suspended by taping the tail 1-2 cm from the tip to prevent grip-based escape attempts. The tape is wrapped securely but not tightly to avoid ischemia. The trial lasts 6 minutes, with the first 1-2 minutes often excluded from analysis to account for initial vigorous struggling. **I**mmobility duration was recorded during the predetermined time (6 minutes) and analyzed with Anymaze software. The state of immobility is defined by the absence of movement (less than 10% area change). The detection of immobility time in these tests serves as a behavioral measure to evaluate the impact of various interventions.

#### Forced swim test (FST)

The FST is used to assess depression-like behavior in rodents by measuring their response to an inescapable water environment. Increased immobility time is interpreted as behavioral despair, while decreased immobility suggests antidepressant-like effects. A transparent Plexiglas cylinder (height: 25-30 cm; diameter: 15-20 cm) filled with 18-20 cm of water (23-25°C) prevents mice from touching the bottom or escaping. Water depth is standardized to ensure mice cannot stabilize by tail-touching. We place the mice individually in a water-filled container for 6 minutes and record their swim and immobility behaviors. The trial duration is 6 minutes, with the first 2 minutes excluded from the analysis to focus on stable behavioral responses, which were analyzed with Anymaze software. We compile the data and calculate the mean immobility time for each experimental group.

#### RNAscope

*Cftr* mRNA expression was assessed in murine brain tissue using a RNAscope Multiplex Fluorescent V2 Assay (Advanced Cell Diagnostics, Bio-Techne, Cat. #323280), following the manufacturer’s protocol. Coronal brain sections (12 μm thick) were collected using a cryostat and stored at −80°C until processing. On the day of the assay, sections were fixed in 4% paraformaldehyde (PFA) in 1X PBS at 4°C for 1 hour, then dehydrated through a graded ethanol series (50%, 70%, and 100%).

To improve tissue permeability, sections were treated with Protease IV at room temperature for 30 minutes. The RNAscope™ Probe—Mm-*Cftr*-C3 (ACDBio, Cat. #483011-C3) was hybridized at 40°C for 2 hours using the HybEZ Oven. Signal amplification was performed according to standard protocol, and *Cftr* transcripts were visualized using TSA Vivid Fluorophore 650 (ACDBio, Cat. #323273), corresponding to the far-red (Cy5) channel. Nuclei were counterstained with DAPI.

Slides were mounted using ProLong™ Gold Antifade Mountant (Thermo Fisher Scientific) to preserve fluorescence. Imaging was performed on four coronal sections spanning the amygdala, hypothalamus, and hippocampus, as described below. Three male and three female mouse brains were assessed correlating to approximately -2.15 mm Bregma.

#### Keyence Imaging

Fluorescent imaging was performed using a Keyence BZ-X700 All-in-One microscope with 10× and 20× objectives. Brain sections containing the amygdala, hypothalamus, and hippocampus were imaged to assess *Cftr* mRNA expression. The far-red channel (TSA Vivid Fluorophore 650, *Cftr* mRNA, displayed as magenta for visualization purposes) was captured using fixed exposure times of 2.5 seconds at 4x, 1.2 seconds at 10×, and 1.5 seconds at 20× to ensure consistency across samples. DAPI exposure was adjusted individually to avoid signal saturation. Images were acquired using Keyence Analyzer software.

#### Image Processing for Visualization

Fluorescent images of *Cftr* mRNA expression in the amygdala, hypothalamus, and hippocampus were imported into ImageJ (FIJI distribution) for visual enhancement and background reduction. To improve edge definition and contrast, the *Unsharp Mask* filter (Process > Filters > Unsharp Mask) was applied using a radius of 1.5 pixels and a mask weight of 0.6. Background signal was removed using the *Subtract Background* function (Process > Subtract Background) with a rolling ball radius of 50 pixels and “Light background” unchecked. These processing steps were applied consistently across all images to facilitate clearer visualization of probe signal while preserving morphological features. Paxino’s and Franklin’s the Mouse Brain in Stereotaxic Coordinates (Plate 49, Bregma -2.15 mm (15)) was used to identify and label brain regions.

#### Drugs

Elexacaftor (cat. # S8851) and ivacaftor (cat. # S1144) were purchased from Selleckchem. Elexacaftor or ivacaftor were separately dissolved in DMSO to prepare 10x stock solutions. Each stock was diluted 1:9 (v/v) with sterile 0.9% saline, vortexed for 30 - 60 seconds and inspected for precipitation. The vehicle control consisted of 10% DMSO in sterile 0.9% saline without drug. Solutions were sterilized by 0.22-µm filtration. An intraperitoneal (IP) injection of elexacaftor (200 mg/kg), ivacaftor (150 mg/kg), or vehicle control were administered to male and female mice, with experimental groups closely matched for sex, age, and weight to ensure scientific rigor. Doses were calculated based on human equivalents using body surface area normalization, as described by Reagan-Shaw et al. (16). An acute treatment paradigm was employed to minimize potential compensatory adaptations associated with repeated dosing. Behavioral assessments were conducted one-hour post-injection.

#### Statistical analysis

One-way analysis of variance (ANOVA) and Tukey’s post hoc multiple comparison tests were used for the statistical comparison. p< 0.05 was considered statistically significant. The graphing and statistical analysis software GraphPad Prism 10 was used to analyze statistical data, which was presented as means ± SEM. In behavioral studies, animals were randomly assigned to experimental groups based solely on their cage numbers. For RNAscope studies, the goal was to confirm presence of Cftr mRNA in both male and female murine brains. We did not perform quantitative analysis.

## Results

### Elexacaftor and ivacaftor induce anxiety-like behaviors

In the OFT **(Fig. 1A)**, a novel environment that elicits both exploratory and anxiety-like responses, mice treated with elexacaftor exhibited significantly reduced locomotor activity compared to the vehicle group **(Fig. 1B)**. Ivacaftor had no effect on locomotion **(Fig. 1B)**. Both elexacaftor- and ivacaftor-treated groups spent less time in the center of the open field, indicative of heightened anxiety-like behavior **(Fig. 1C)**.

**Fig. 1.**
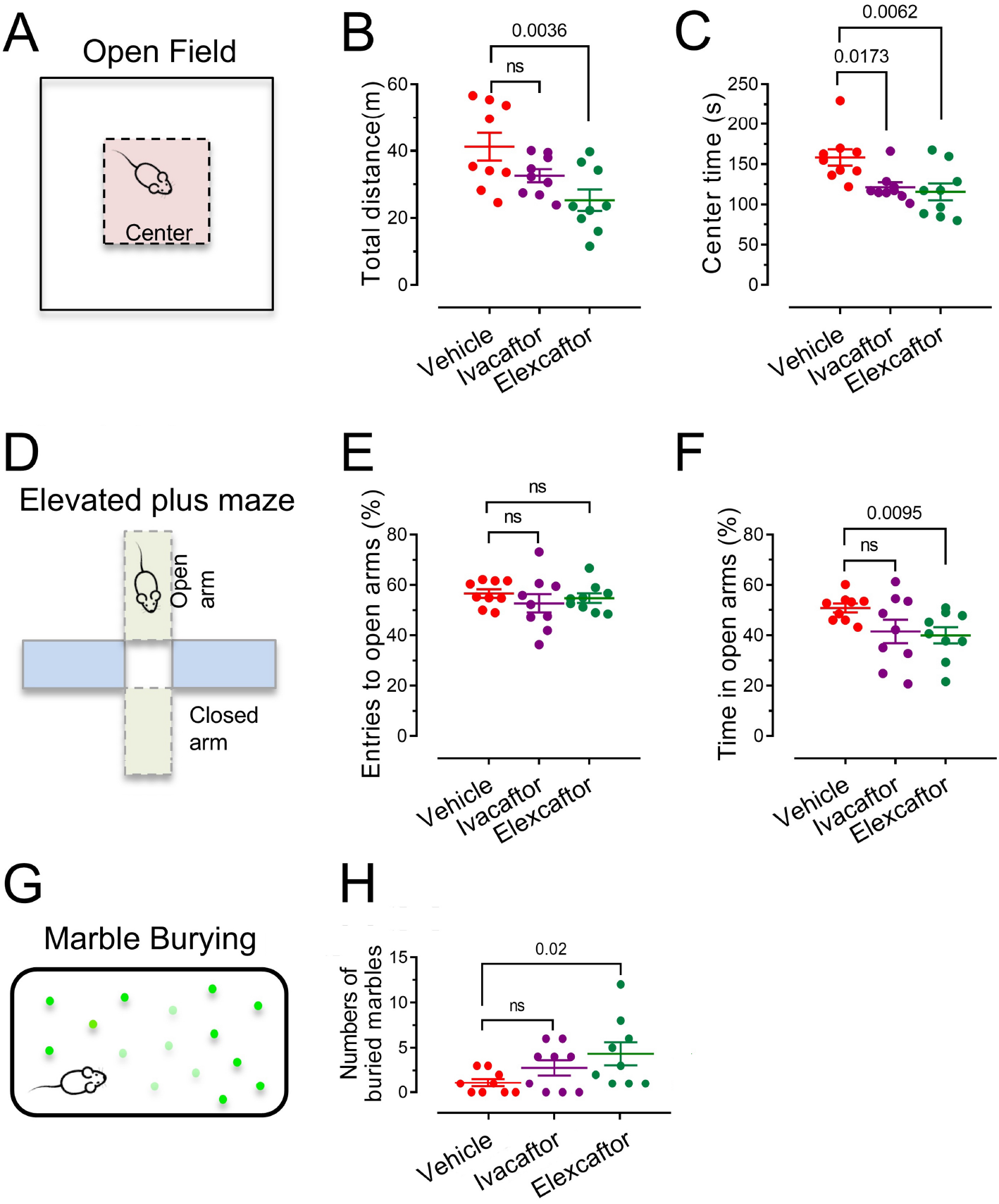
Effects of elexacaftor and ivacaftor on anxiety-like behaviors. **(A)** Schematic of the open field test. **(B)** Total travel time and **(C)** the time spend in the center area. **(D)** Schematic of the elevated plus maze test. **(E)** Entries to open arms and **(F)** the time spend in open arms. **(G)** Schematic of the marble burying test and **(H)** the numbers of the buried marbles. Data are shown as mean ± SEM. Student’s t-test. ns, non-significant. *p* values are shown in each panel. Vehicle, n = 4 females and 5 males; Ivacaftor, n = 4 females and 5 males; Elexacaftor n = 4 females and 5 males.

In the EPMT **(Fig. 1D)**, reduced time in open arms reflects increased anxiety. While neither elexacaftor nor ivacaftor altered the number of open-arm entries **(Fig. 1E)**, elexacaftor significantly decreased the time spent in the open arms compared to the vehicle group **(Fig. 1F)**, suggesting anxiety-like behavior. This effect was not observed for ivacaftor.

In the MBT, elexacaftor-treated mice displayed a significant increase in marble burying behavior relative to the vehicle group (**Fig. 1H**), further supporting an anxiogenic effect. Ivacaftor did not significantly alter marble burying behavior.

Collectively, these findings demonstrate that acute administration of elexacaftor consistently induced anxiety-like behavior across multiple behavioral paradigms, while ivacaftor exhibits a more limited effect, primarily influencing center time in the OFT. These results suggest that elexacaftor, and to a lesser extent ivacaftor, may modulate neural circuits, potentially involving the amygdala, to promote anxiogenic responses in mice

### Ivacaftor induces depressive-like behaviors

In the NSFT **(Fig. 2A)**, which leverages the suppression of feeding behavior in a novel environment as an indicator of depressive-like behavior, ivacaftor significantly increased the latency to initiate feeding compared to the vehicle group **(Fig. 2B)**. Additionally, ivacaftor-treated mice exhibited a marked reduction in the time spent sniffing the food pellet **(Fig. 2C)**, further suggesting an enhanced depressive-like phenotype. In contrast, elexacaftor did not significantly alter either latency to feed or sniffing behavior, indicating a lack of effect on NSFT performance. These findings suggest that ivacaftor, but not elexacaftor, promoted behaviors consistent with heightened depressive-like states in this paradigm.

**Fig. 2.**
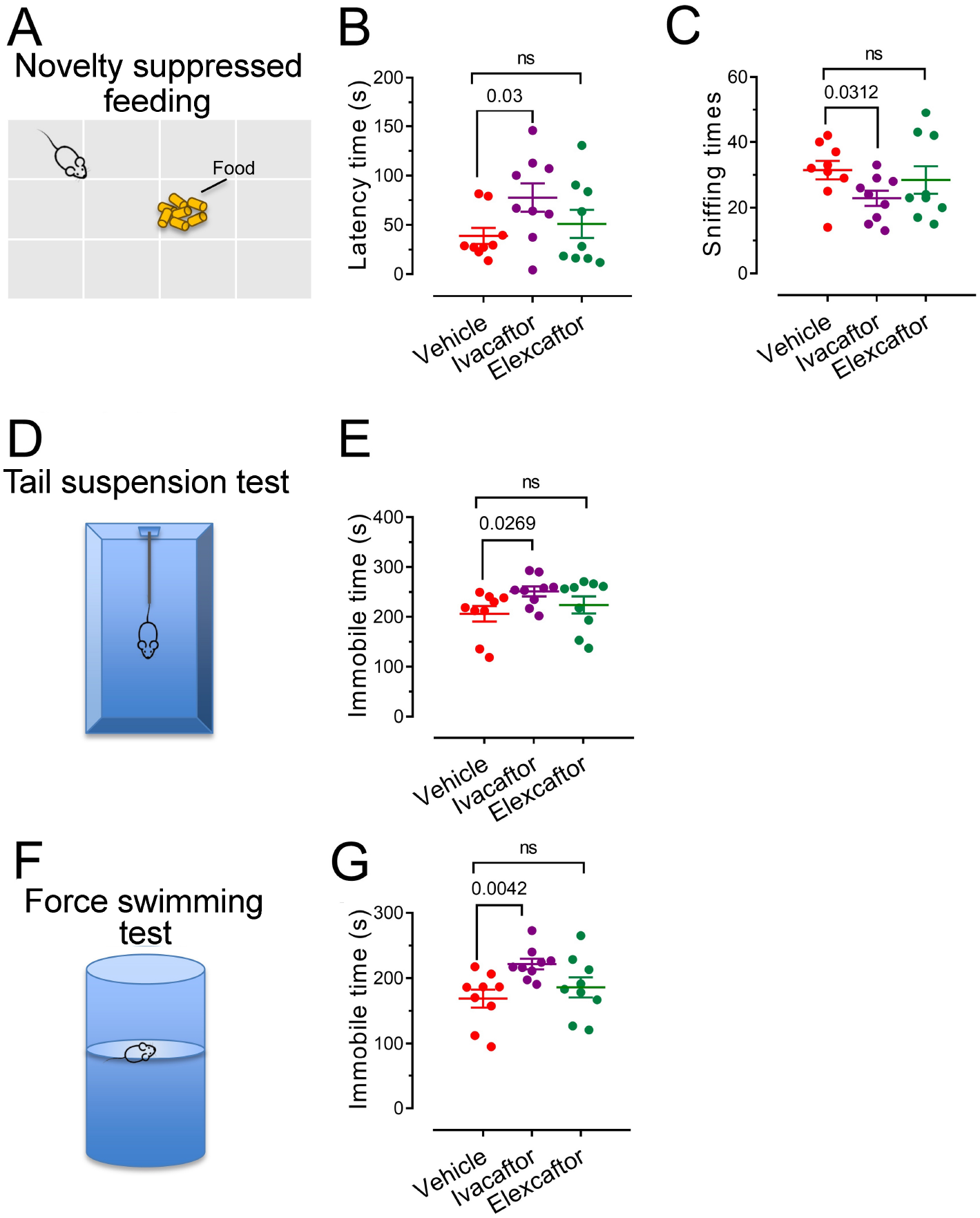
Effects of elexacaftor and ivacaftor on depressive-like behaviors. **(A)**, schematic of the Novelty suppressed feeding test, **(B)** results of latency time and **(C)** sniffing time. **(D)**, schematic of the tail suspension test and **(E)** the results of immobile time. **(F)**, schematic of the force swimming test and **(G)** the results of immobile time. Data are shown as mean ± SEM. Student’s t-test. ns, non-significant. *p* values are shown in each panel. For all panels, Vehicle, n = 4 females and 5 males; Ivacaftor, n = 4 females and 5 males; Elexacaftor n = 4 females and 5 males.

The TST **(Fig. 2D)**, a widely used assay to measure stress-induced immobility as a proxy for behavioral despair, revealed similar trends. Mice treated with ivacaftor displayed significantly increased immobility time compared to the vehicle group **(Fig. 2E)**, consistent with a depressive-like response akin to helplessness. Elexacaftor-treated mice, however, showed no significant difference in immobility time relative to controls, reinforcing the specificity of ivacaftor’s effect on depressive-like behavior. The FST **(Fig. 2F)**, another robust model for evaluating despair-like behavior, corroborated these results. Ivacaftor-treated mice exhibited prolonged immobility times compared to the vehicle group, while elexacaftor had no significant effect on FST performance **(Fig. 2G)**. These consistent outcomes across the TST and FST underscore ivacaftor’s propensity to induce depressive-like behaviors.

### CFTR mRNA is found in brain regions that regulate mood, including the amygdala and hippocampus

McGrath and colleagues were the first to identify CFTR expression in the nervous system (17) and subsequent studies by Mulberg demonstrated its widespread distribution in the rat brain (18, 19). Consistent with these findings, we confirmed broad *Cftr* mRNA expression in the mouse brain (**Fig. 3A**), and specifically in regions that regulate mood (**Fig. 3 A-D**), including the hippocampus (**Fig. 3B**), the amygdala (**Fig. 3C**), and hypothalamus (**Fig. 3D**). These results support earlier observations and suggest that the neurobehavioral effects of elexacaftor and ivacaftor may arise from direct modulation of CFTR within the brain.

**Fig. 3.**
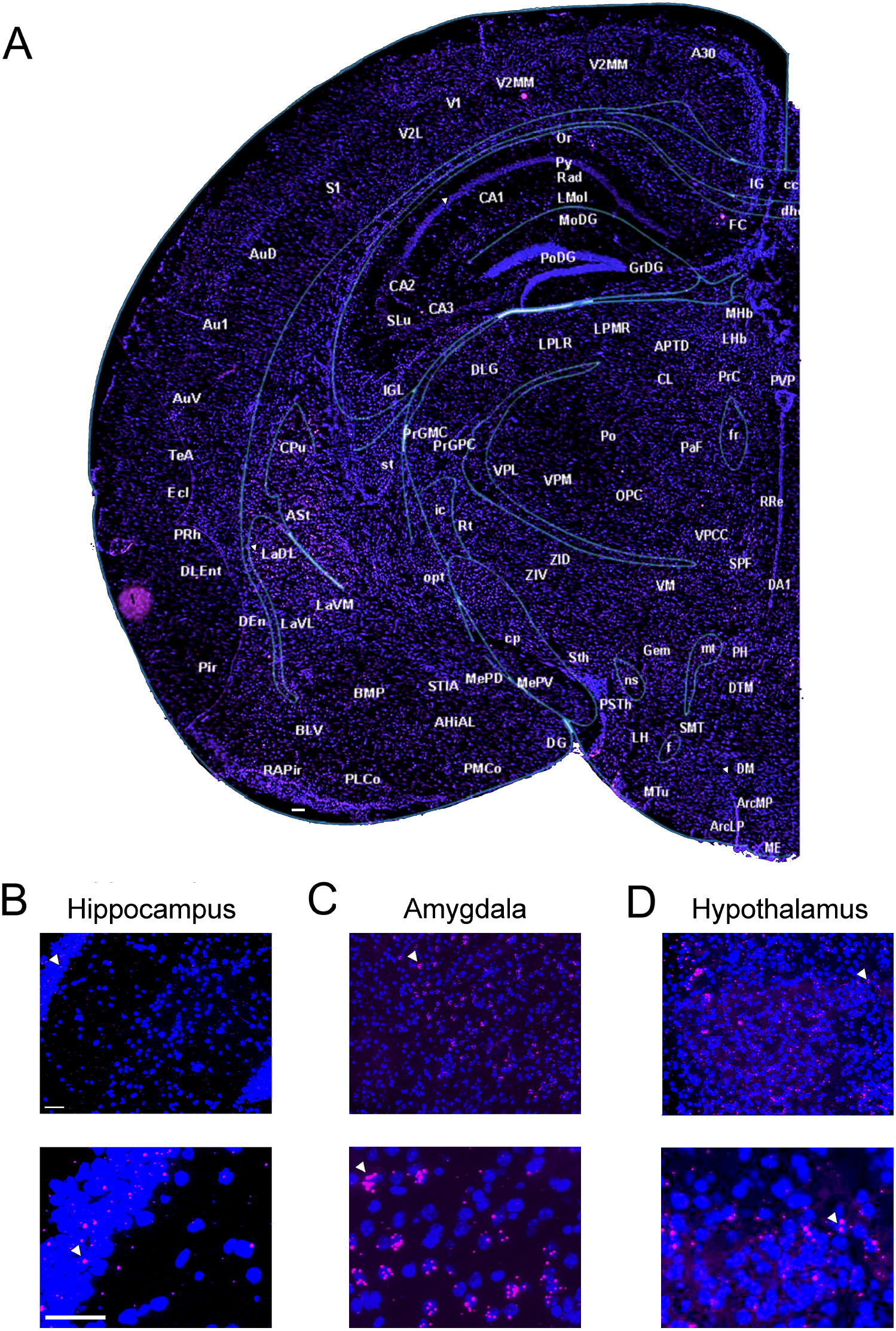
*Cftr* mRNA in the murine brain. **(A) Representative** stitched image showing half hemisphere of murine brain with widespread *Cftr* mRNA expression. RNAscope was used to assess Cftr mRNA (illustrated by pink/purple fluorescence). Dapi to stain cell nuclei are shown in blue. Scale bar is 100 μm. Labels and abbreviations correspond to those provided by (15). Small arrows in the hippocampus, amygdala, and hypothalamus represent same cells in subsequent higher power magnification panels. **(B)** Hippocampus. Bottom panel shows more magnified view. Scale bars in both panels are 50 μm. Arrow indicates same mRNA molecule(s) in each panel and location as shown in A. **(C)** Amygdala. We focused on the lateral amygdala, which showed qualitatively robust mRNA expression. Bottom panel shows more magnified view. Scale bars in both panels are 50 μm. Arrow indicates same mRNA molecule(s) in each panel and location as shown in A. **(D)**, Hypothalamus. Bottom panel shows more magnified view. Scale bars in both panels are 50 μm. Arrow indicates same mRNA molecule(s) in each panel and location as shown in A. A total of 3 male and 3 female brains were examined. Quantitative assessment was not performed.

## Discussion

The present study provides novel evidence that CFTR modulators, elexacaftor and ivacaftor, induce anxiety- and depressive-like behaviors in mice, potentially through direct actions on CFTR-expressing regions of the brain. These findings align with clinical observations of worsening anxiety and depression in some people with CF treated with Trikafta. Our data underscore the critical need to investigate the neural mechanisms underlying the psychiatric side effects of CFTR modulators and consider whether they are due to direct modulation of CFTR in the brain or other mechanisms.

The clinical benefits of CFTR modulators, particularly Trikafta, in improving lung function and survival in CF patients are well-documented (20). However, emerging reports highlight a paradoxical deterioration in mental health among some people, with anxiety and depression noted as adverse effects (see (10) for recent review). Our preclinical findings mirror these clinical observations: acute administration of elexacaftor and ivacaftor in mice induced distinct behavioral phenotypes - elexacaftor heightened anxiety-like behaviors (reduced center time in OFT, decreased open-arm exploration in EPMT, and increased marble burying), while ivacaftor exacerbated depressive-like behaviors (prolonged latency in NSFT, increased immobility in TST and FST). These results suggest that the two drugs may differentially modulate neural circuits governing emotional states. Importantly, tezacaftor, the third component of Trikafta, was excluded from this study due to its low blood-brain barrier penetrance, narrowing the focus to elexacaftor and ivacaftor as potential CNS-active agents.

The distinct neurobehavioral profiles associated with ivacaftor and elexacaftor - ivacaftor primarily inducing depressive-like behaviors and elexacaftor primarily inducing anxiety-like behaviors - raise important questions about their underlying mechanisms. Ivacaftor functions as a CFTR potentiator, while elexacaftor is primarily a corrector with some potentiator activity (21). One possibility is that ivacaftor’s potentiation of existing CFTR disrupts normal chloride homeostasis in the central nervous system, leading to an imbalance between excitatory and inhibitory neurotransmission and a net depressive effect. Conversely, elexacaftor may enhance CFTR trafficking and function in a manner resembling CFTR overexpression, potentially resulting in increased chloride conductance and a net anxiogenic outcome. Within this framework, potentiation of existing CFTR may contribute to depressive-like behaviors, whereas increased CFTR expression may promote anxiety-like behaviors. However, to our knowledge, there is currently no direct evidence that elexacaftor increases wild-type CFTR protein expression. An alternative explanation is that the observed neurobehavioral effects are CFTR-independent, especially given that ivacaftor has known off-target activity at several neurotransmitter receptors (22).

While our findings are compelling, several limitations warrant cautious interpretation. First, this study employed acute drug administration in wild-type mice; thus, the neurobehavioral effects observed may not directly translate to mice with CFTR mutations. The interaction between CFTR dysfunction, modulator therapy, and mental health is likely more complex in individuals with cystic fibrosis than in healthy animal models. Second, our dosing paradigm does not replicate the chronic treatment regimens typically used in people with CF. We selected an acute dosing strategy to isolate immediate neurobehavioral effects, minimizing confounding influences from neuroplastic or adaptive changes that may arise with prolonged exposure. Third, although intraperitoneal injection is standard in preclinical research, it differs from the oral administration used in humans and may alter pharmacokinetics and central nervous system drug exposure. Finally, while our behavioral assays are validated proxies for anxiety- and depression-like phenotypes, they offer indirect measures. Integrating complementary approaches, such as electrophysiological recordings or molecular profiling of stress-related pathways (e.g., corticotropin-releasing hormone, serotonin), would enhance mechanistic insights.

Future studies should prioritize several avenues. First, replicating these experiments in CF animal models (e.g., ΔF508 mutants) will clarify the drugs’ neurobehavioral effects. Second, chronic dosing studies are essential to determine if tolerance develops or if side effects worsen over time or change over time. Third, mechanistic investigations should explore how elexacaftor and ivacaftor alter synaptic function in the brain, with areas like the amygdala and hippocampus emphasized. For example, CFTR activation might impair long-term potentiation (LTP) or promote excitatory/inhibitory imbalance, as seen in mood disorders. Techniques such as patch-clamp electrophysiology or calcium imaging in amygdala slices could elucidate these processes. Finally, translational research should identify biomarkers (e.g., amygdala or hippocampal activity on fMRI, chloride levels in cerebrospinal fluid) to predict and monitor mental health outcomes in people with CF on modulator therapies.

In summary, this study provides preclinical evidence that elexacaftor and ivacaftor, key components of Trikafta, induce anxiety- and depressive-like behaviors in mice. While our findings do not conclusively determine whether these effects result from direct modulation of CFTR in the brain, the presence of *Cftr* mRNA expression in limbic regions supports this possibility. Further studies are needed to elucidate the underlying mechanisms, including the potential role of central CFTR modulation, and to assess the translational relevance of these findings to individuals with CF.

## Author contributions

J.D. and L.R.R. conceived the project. J.D., L.R.R., H.L., A.F. and Q.G. designed the experiments. Q.G. and H.L. performed behavior experiments and data analysis. A.F. performed the RNAscope experiments and data analysis. L.R.R., J.D., H.L., A.F. and Q.G. wrote the manuscript. All authors reviewed and edited the manuscript.

## Acknowledgments

J.D. is supported by the National Institutes of Mental Health (R01MH135862) and the Cystic Fibrosis Foundation (LI21G0). L.R.R is supported by the National Heart Lung and Blood Institute (R01HL152101) and the Cystic Fibrosis Foundation (REZNI23Y3). H.L is supported by the National Institutes Neurological Disorders and Stroke (R01NS118197). The authors thank Shanil Amin for technical assistance.

## Conflict-of-interest

The authors have declared that no conflict of interest exists.

## Notes

### Competing Interest Statement

The authors have declared no competing interest.

## References

1. S. M. Hoy, Elexacaftor/Ivacaftor/Tezacaftor: First Approval. Drugs 79, 2001–2007 (2019).

2. M. E. Condren, M. D. Bradshaw, Ivacaftor: a novel gene-based therapeutic approach for cystic fibrosis. J Pediatr Pharmacol Ther 18, 8–13 (2013).

3. E. D. Deeks, Lumacaftor/Ivacaftor: A Review in Cystic Fibrosis. Drugs 76, 1191–1201 (2016).

4. T. Havermans, L. Willem, Prevention of anxiety and depression in cystic fibrosis. Curr Opin Pulm Med 25, 654–659 (2019).

5. I. Baiardini, G. Steinhilber, D. I. M. FF. Braido, P. Solidoro, Anxiety and depression in cystic fibrosis. Minerva Med 106, 1–8 (2015).

6. K. A. Riekert, S. J. Bartlett, M. P. Boyle, J. A. Krishnan, C. S. Rand, The association between depression, lung function, and health-related quality of life among adults with cystic fibrosis. Chest 132, 231–237 (2007).

7. C. Snell, S. Fernandes, I. S. Bujoreanu, G. Garcia, Depression, illness severity, and healthcare utilization in cystic fibrosis. Pediatr Pulmonol 49, 1177–1181 (2014).

8. A. M. Yohannes, T. G. Willgoss, F. A. Fatoye, M. D. Dip, K. Webb, Relationship between anxiety, depression, and quality of life in adult patients with cystic fibrosis. Respir Care 57, 550–556 (2012).

9. T. Havermans, K. Colpaert, L. J. Dupont, Quality of life in patients with Cystic Fibrosis: association with anxiety and depression. J Cyst Fibros 7, 581–584 (2008).

10. C. J. Bathgate et al., Elexacaftor/tezacaftor/ivacaftor and mental health: A workshop report from the Cystic Fibrosis Foundation’s Prioritizing Research in Mental Health working group. J Cyst Fibros 24, 301–309 (2025).

11. L. R. Reznikov, Cystic Fibrosis and the Nervous System. Chest 151, 1147–1155 (2017).

12. W. Haefely, Benzodiazepine interactions with GABA receptors. Neurosci Lett 47, 201–206 (1984).

13. J. L. Barker, D. G. Owen, M. Segal, GABA actions on the excitability of cultured CNS neurons. Neurosci Lett 47, 313–318 (1984).

14. A. A. Walf, C. A. Frye, The use of the elevated plus maze as an assay of anxiety-related behavior in rodents. Nat Protoc 2, 322–328 (2007).

15. K. B. J. Franklin, & Paxinos, G., The mouse brain in stereotaxic coordinates (4th ed.). (Academic Press., 2012).

16. S. Reagan-Shaw, M. Nihal, N. Ahmad, Dose translation from animal to human studies revisited. FASEB J 22, 659–661 (2008).

17. S. A. McGrath, A. Basu, P. L. Zeitlin, Cystic fibrosis gene and protein expression during fetal lung development. Am J Respir Cell Mol Biol 8, 201–208 (1993).

18. A. E. Mulberg et al., Cystic fibrosis transmembrane conductance regulator protein expression in brain. Neuroreport 5, 1684–1688 (1994).

19. A. E. Mulberg et al., Expression and localization of the cystic fibrosis transmembrane conductance regulator mRNA and its protein in rat brain. J Clin Invest 96, 646–652 (1995).

20. M. Shteinberg, J. L. Taylor-Cousar, Impact of CFTR modulator use on outcomes in people with severe cystic fibrosis lung disease. Eur Respir Rev 29 (2020).

21. O. Laselva et al., Rescue of multiple class II CFTR mutations by elexacaftor+tezacaftor+ivacaftor mediated in part by the dual activities of elexacaftor as both corrector and potentiator. Eur Respir J 57 (2021).

22. E. K. Schneider et al., The potentially beneficial central nervous system activity profile of ivacaftor and its metabolites. ERJ Open Res 4 (2018).

